# rBahadur: Efficient simulation of high-dimensional genotype data with global dependence structures

**DOI:** 10.1101/2022.10.13.512132

**Authors:** Richard Border, Osman Asif Malik

## Abstract

Existing methods for generating synthetic genotype data are ill-suited for replicating the effects of assortative mating (AM). We propose `rb_dplr`, a novel and computationally efficient algorithm for generating high-dimensional binary random variates that effectively recapitulates AM-induced genetic architectures using the Bahadur order-2 approximation of the multivariate Bernoulli distribution.

**Availability and implementation:** The `rBahadur` R library is available at https://github.com/rborder/rBahadur.

---

The simulation of realistic genotype / phenotype data is a fundamental tool in statistical genetics and is essential for the development of robust statistical methods for the analysis of genome-wide data. As such, much prior effort has focused on generating synthetic data that recapitulate salient characteristics of genetic marker data, including local linkage disequilibrium (LD) structure induced by variable recombination rates [1, 2, 3], relationships between local LD structure and allelic effects [4], and the many consequences of drift, admixture, and geographic stratification [5, 6].

Despite these advances, existing methods are ill-suited for generating synthetic genotype / phenotype data reflecting the consequences of recent assortative mating (AM); in contrast to the effects of recombination, which yields banded covariance structures, AM induces dense covariance structures reflecting sign-consistent dependence among all causal variants across the genome [7, 8]. On the other hand, there is substantial recent evidence that AM is widespread [9, 8, 10] and complicates the interpretation of many commonly applied methods in statistical genetics, including heritability estimation [11], genetic correlation estimation [8, 12], and Mendelian randomization [13]. Efficient simulation methods for generating high-dimensional genotype / phenotype data congruent with the consequences of AM will be critical to the development of robust analytic tools.

SNP haplotypes subject to AM-induced long-range dependencies can be represented mathematically by the *m*-dimensional multivariate Bernoulli (MVB) distribution [14], which is challenging to sample from; existing methods require *O*(*m*^2^) or more operations to draw a vector of *m* haplotypes (see Figure 1), which becomes infeasible in high dimensions. As such, existing methods for synthesizing AM-consistent marker data at scale obviate this problem by generating genotype / phenotype data assuming random mating and subsequently proceeding through multiple generations of forward simulation, complete with mating and meioses [2, 11, 8]. However, these methods require repeatedly shuffling the elements of large arrays and require simulating the genotypes of a large number of individuals to obtain a relatively small sample of unrelated individuals, making them cumbersome in the context of methods development.

**Figure 1:**
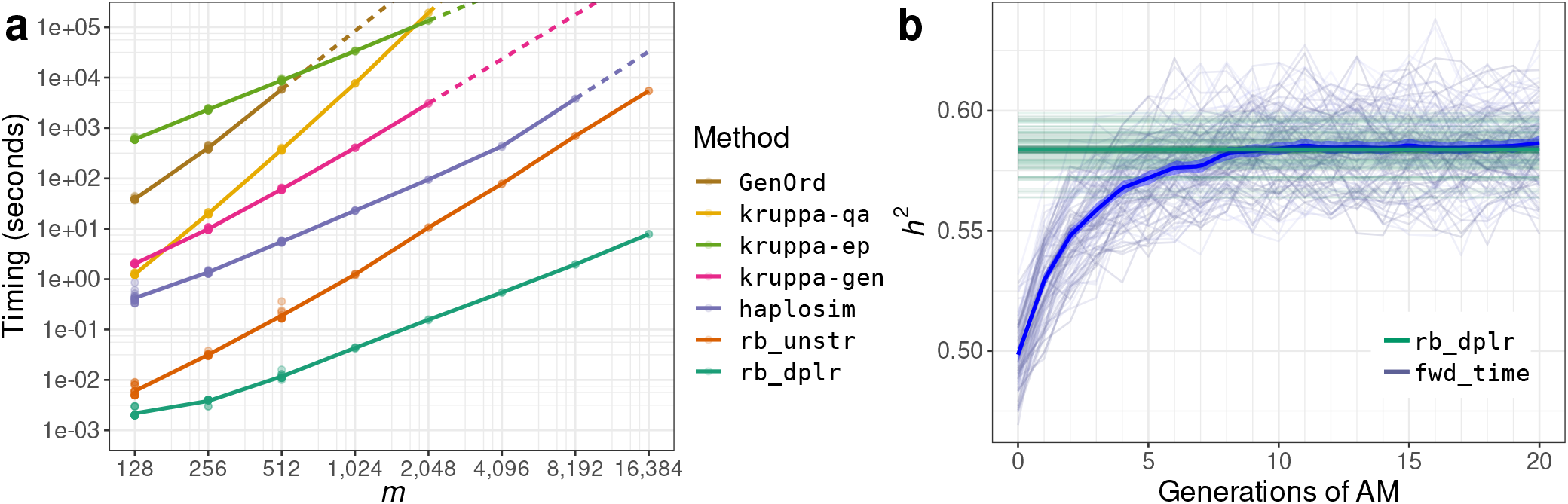
**(a)** Cross-method comparison of single-threaded wall time for generating *m*/4 samples from the m-dimensional MVB distribution on a log-log scale. The proposed `rb_dplr` algorithm scales linearly in sample size and problem dimension. Both `rb_dplr` and the unstructured variant `rb_unstr` outperform existing methods including haplosim [3], three methods implemented by Kruppa et al. (`kruppa-*`) [1], and `GenOrd` [16]. Solid lines reflect linear splines with fixed knots fitted to numerical experiment results and dashed lines reflect extrapolations. **(b)** Drawing genotypes directly from their equilibrium distribution under AM. Comparison of heritabilities in synthetic genotype / phenotype data generated using `rb_dplr` to sample from the appropriate MVB distribution, versus the forward-time approach of [8]. Best-fit lines and standard-errors summarize variation across 100 replicates with 2000 haploid causal variants for 8000 individuals, for phenotypes with panmictic heritability cross-mate phenotypic correlation both fixed to 0.5.

In the current manuscript, we introduce a novel collection of efficient methods for directly sampling highdimensional MVB random variables satisfying particular moment conditions (i.e., admitting a Bahadur order-2 representation; [15]). In particular, we propose the `rb_dplr` algorithm, which exploits the diagonal-plus-low-rank correlation structure induced by AM to generate MVB samples using only *O*(*m*) operations. We then present numerical experiments demonstrating that the proposed methods outperform existing direct sampling methods and verify that they faithfully represent the effects of AM by comparing results to forward-time simulations. We provide these methods, together with a collection of utilities for characterizing the equilibrium distribution of haplotypes under AM, in `rBahadur`, an open-source library for the R programming language.

## Method

Suppose *X*_1_, …, *X_m_* are Bernoulli random variables with means 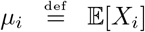. When the variables are independent, the distribution of (*X*_1_, …, *X_m_*) is simply 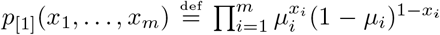, but this is not the case in general. Bahadur [15] showed that the distribution of an MVB takes the form

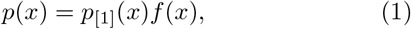

where *x* = (*x*_1_, …, *x_m_*). The function *f* in (1) is defined as

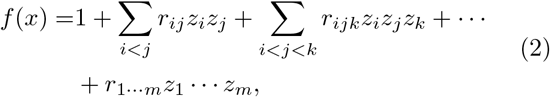

where 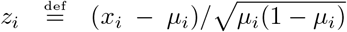 and 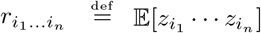. The means (*μ_i_*) and mixed moments (*r_ij_*), (*r_ijk_*), and so forth, characterize the MVB and are comprised of 2^*m*^ – 1 parameters. This exponential dependence on *m* makes working with general MVBs challenging.

We consider Bahadur order-2 approximations to the distribution in (1); i.e., we assume that *r*_*i*_1_…*i_n_*_ = 0 for *n* ≥ 3. Thus, the order-2 MVB distribution is fully characterized by its means (*μ_i_*) and correlations (*r_ij_*). `rBahadur` provides two methods for sampling from this distribution. The first algorithm (`rb_unstr`) can handle generic correlation matrices and requires *O*(*m*^2^) operations to sample from the m-dimensional MVB. The second algorithm (`rb_dplr`) is developed specifically for the case when the correlation matrix is diagonal-plus-low-rank (DPLR; i.e., (*r_ij_*) = *D* + *UU^T^* where *D* is diagonal and *U* is *m* × *c* for some *c* ≪ *m*) and requires *O*(*mc*) operations. Both methods sample the entries of the random vector sequentially: First, a realization of *X*_1_ is drawn. Then, subsequent variables *X_n_* are drawn *conditionally* on the realization of the previously drawn variables *X*_1_, …, *X*_*n*–1_ for 2 ≤ *n* ≤ *m*. For further details, see Supplementary Materials S1.

### Genotype simulation under assortment via rb_dplr

Let *X* denote the standardized genotypes at *m* haploid SNP loci with allele frequencies *μ* and standardized effects *β*. Consider an additive polygenic phenotype *y* with 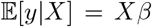 and panmictic heritability 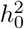. The equilibrium joint distribution of *X* and *β* under primary-phenotypic AM will be *p*(*X, β*) = *p*(*X* |*β*)*p*(*β*) where *p*(*X*|*β*) is a MVB distribution with mean vector *μ* and correlation

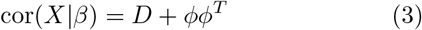

where the vector *ϕ* and the diagonal matrix *D* are known functions of *μ*, *β*, 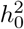 and the cross-mate correlation *r* (Supplementary Materials S2.1). Assuming causal variants are not in strong local LD with one another at panmixis, *p*(*X*|*β*) is well approximated by the corresponding DPLR Bahadur order-2 distribution, which is determined by only *O*(*m*) parameters and can be efficiently sampled using `rb_dplr`. We provide a utility for sampling from this distribution via the `am_corr_struct` function.

## Numerical experiments

Figure 1a compares the time required to generate *m* binary haplotypes for *n* = *m*/4 individuals under the model described above, with 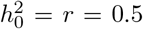, across existing and proposed methods. Both `rb_dplr` and `rb_unstr` outperform existing methods, with `rb_dplr` evidencing *O*(*mn*) time complexity congruent with theoretical expectations. Figure 1b compares heritabilities associated with genotype / phenotype data as generated with `rb_dplr` under the model (3) versus those achieved after of up to 20 generations of the corresponding forward-time procedure, as implemented in [8], across 100 replicates (comparing `rb_dplr` *h*^2^ to mean *h*^2^ values across forward-time generations 16-20, Welch’s *t*(99) = 0.84, *p* = 0.40). Here, `rb_dplr` provides a direct and efficient alternative to forward time approaches that can be readily incorporated into sensitivity analysis and methods development pipelines.

## Discussion

Given that AM is both widespread [11, 10] and consequential for the interpretation of marker-based estimators [8, 11, 13], it is crucial that statistical geneticists are able to perturb random-mating assumptions when developing and evaluating methods. To this end, we have developed the `rBahadur` library to efficiently sample haploid causal variants under AM-induced genetic architectures. The software is open-source and freely available at https://github.com/rborder/rBahadur.

Our approach is limited by the requirement that the target distribution admits a second-order Bahadur approximation, limiting applications to complex correlation structures involving both strong local LD and AM-induced global dependence. We address this by ensuring the simulation functions in `rBahadur` fail transparently in such cases and describe how `rBahadur` can be used in conjunction with reference haplotypes to overcome this limitation in Supplementary Materials S2.2.

