## Supplementary Materials for "rBahadur: Efficient simulation of high-dimensional genotype data with global dependence structures"

Richard Border[†‡1] and Osman Asif Malik[†2]

[1]Departments of Neurology and Computer Science, University of California, Los Angeles
[2]Applied Mathematics & Computational Research Division, Lawrence Berkeley National Laboratory

#### S1 Methods for sampling MVB random variates

We consider the problem of drawing the realization of $m$ random variables $X_1, X_2, \ldots, X_m$ from the multivariate Bernoulli (MVB) distribution. For $m$ such random variables, there are $2^m$ possible outcomes. Following [1], for $k \in [2^m] \stackrel{\text{def}}{=} \{1, 2, \ldots, 2^m\}$, let $(k_1, k_2, \ldots, k_m)$ be the binary expansion of $k$ defined via

$$k = 1 + \sum_{n=1}^{m} k_n 2^{n-1}, \tag{1}$$

where each $k_n \in \{0, 1\}$. Let $\mathbf{p}^{(m)} \in \mathbb{R}^{2^m}$ be a vector of probabilities defined as

$$\mathbf{p}_k^{(m)} \stackrel{\text{def}}{=} \mathbb{P}\{X_1 = k_1, X_2 = k_2, \ldots, X_m = k_m\}.$$

Let $\boldsymbol{\sigma}^{(m)} \in \mathbb{R}^{2^m}$ be a vector of (central) moments, defined elementwise via

$$\boldsymbol{\sigma}_k^{(m)} \stackrel{\text{def}}{=} \mathbb{E}\Big(\prod_{n=1}^{m} (X_n - \mu_n)^{k_n}\Big),$$

where $\mu_n \stackrel{\text{def}}{=} \mathbb{E}[X_n]$ is the $n$th marginal probability. We assume each $\mu_n \in (0, 1)$ as other the corresponding random variable is simply constant. Note that $\boldsymbol{\sigma}_1^{(m)} = 1$, and $\boldsymbol{\sigma}_k^{(m)} = 0$ for all $k = (k_1, \ldots, k_m)$ such that $\sum_n k_n = 1$ (i.e., all first order central moments are zero). For an index $k \in [2^n]$ with binary expansion $(k_1, \ldots, k_n)$, we use will use the notations $\boldsymbol{\sigma}_k^{(n)}$ and $\mathbf{p}_k^{(n)}$ interchangeably with $\boldsymbol{\sigma}_{k_1 \cdots k_n}^{(n)}$ and $\mathbf{p}_{k_1 \cdots k_n}^{(n)}$.

As discussed in the main paper, we only consider MVB distributions of Bahadur order-2. This is made precise by the following assumption.

**Assumption S1.1.** We assume that third order central moments and higher in $\boldsymbol{\sigma}^{(m)}$ are zero.

†Authors contributed equally.

The second order moments in $\boldsymbol{\sigma}^{(m)}$ are precisely those on positions $k = (k_1, \dots, k_m)$ with $\sum_n k_n = 2$. Table 1 shows an example of $\boldsymbol{\sigma}^{(m)}$ in the case $m = 4$, where $\sigma_{ij}$ denotes the second order central moment involving $X_i$ and $X_j$.

For a vector $\mathbf{x} \in \mathbb{R}^m$ and integers $a$ and $b$ satisfying $1 \leq a < b \leq m$, let $\mathbf{x}_{a:b}$ denote the subvector

$$\mathbf{x}_{a:b} \stackrel{\text{def}}{=} \begin{bmatrix} \mathbf{x}_a \\ \mathbf{x}_{a+1} \\ \vdots \\ \mathbf{x}_b \end{bmatrix}.$$

Notice that $\boldsymbol{\sigma}^{(4)}_{1:8} = \boldsymbol{\sigma}^{(3)}$. This is true in general: $\boldsymbol{\sigma}^{(n+1)}_{1:2^n} = \boldsymbol{\sigma}^{(n)}$ for $n \geq 1$. Defining $\boldsymbol{\gamma}^{(n)} \stackrel{\text{def}}{=} \boldsymbol{\sigma}^{(n+1)}_{2^n+1:2^{n+1}}$, we have

$$\boldsymbol{\sigma}^{(n+1)} = \begin{bmatrix} \boldsymbol{\sigma}^{(n)} \\ \boldsymbol{\gamma}^{(n)} \end{bmatrix}, \quad n \geq 1. \tag{2}$$

We will use $\tilde{\boldsymbol{\gamma}}^{(n)} \in \mathbb{R}^n$ to denote the vector containing the $n$ potentially nonzero elements in $\boldsymbol{\gamma}^{(n)}$. More precisely,

$$\tilde{\boldsymbol{\gamma}}^{(n)} \stackrel{\text{def}}{=} \begin{bmatrix} \sigma_{1(n+1)} \\ \sigma_{2(n+1)} \\ \vdots \\ \sigma_{n(n+1)} \end{bmatrix}. \tag{3}$$

For each $n \in [N]$, let $\mathbf{B}^{(n)} \in \mathbb{R}^{2\times 2}$ be defined as

$$\mathbf{B}^{(n)} \stackrel{\text{def}}{=} \begin{bmatrix} 1-\mu_n & -1 \\ \mu_n & 1 \end{bmatrix}.$$

From Theorem 1 in [1], we have

$$\mathbf{p}^{(m)} = (\mathbf{B}^{(m)} \otimes \mathbf{B}^{(m-1)} \otimes \cdots \otimes \mathbf{B}^{(1)}) \boldsymbol{\sigma}^{(m)}, \tag{4}$$

where $\otimes$ denotes the matrix Kronecker product.

In the following, we will use the notation

$$p_{k_1\cdots k_n} \stackrel{\text{def}}{=} \mathbb{P}\{X_1 = k_1, \dots, X_n = k_n\},$$
$$p_{k_n|k_1\cdots k_{n-1}} \stackrel{\text{def}}{=} \mathbb{P}\{X_n = k_n \mid X_1 = k_1, \dots, X_{n-1} = k_{n-1}\}.$$

To simplify notation, we will use zero based indexing for the matrices $\mathbf{B}^{(n)}$, so that, e.g., $\mathbf{B}^{(n)}_{10} = \mu_n$ and $\mathbf{B}^{(n)}_{01} = -1$.

**Lemma S1.2.** *Assume $p_{k_1\cdots k_n} > 0$. The conditional probability $p_{k_n|k_1\cdots k_{n-1}}$ satisfies*

$$p_{k_n|k_1\cdots k_{n-1}} = \mathbf{B}^{(n)}_{k_n 0} + \mathbf{B}^{(n)}_{k_n 1} \sum_{j_1,\dots,j_{n-1}} \frac{\mathbf{B}^{(n-1)}_{k_{n-1}j_{n-1}} \cdots \mathbf{B}^{(1)}_{k_1 j_1}}{p_{k_1\cdots k_{n-1}}} \boldsymbol{\gamma}^{(n-1)}_{j_1\cdots j_{n-1}}, \tag{5}$$

*where the sum is over indices $(j_1, \dots, j_{n-1}) \in \{0,1\}^{n-1}$.*

*Proof.* From (4), it follows that

$$p_{k_1\cdots k_n} = \sum_{j_1,\dots,j_n} \mathbf{B}^{(n)}_{k_n j_n} \cdots \mathbf{B}^{(1)}_{k_1 j_1} \boldsymbol{\sigma}^{(n)}_{j_1\cdots j_n},$$

| | Binary expansion | | | | |
|---|---|---|---|---|---|
| $k$ | $k_1$ | $k_2$ | $k_3$ | $k_4$ | $\boldsymbol{\sigma}_k^{(4)}$ |
| 1 | 0 | 0 | 0 | 0 | 1 |
| 2 | 1 | 0 | 0 | 0 | 0 |
| 3 | 0 | 1 | 0 | 0 | 0 |
| 4 | 1 | 1 | 0 | 0 | $\sigma_{12}$ |
| 5 | 0 | 0 | 1 | 0 | 0 |
| 6 | 1 | 0 | 1 | 0 | $\sigma_{13}$ |
| 7 | 0 | 1 | 1 | 0 | $\sigma_{23}$ |
| 8 | 1 | 1 | 1 | 0 | 0 |
| 9 | 0 | 0 | 0 | 1 | 0 |
| 10 | 1 | 0 | 0 | 1 | $\sigma_{14}$ |
| 11 | 0 | 1 | 0 | 1 | $\sigma_{24}$ |
| 12 | 1 | 1 | 0 | 1 | 0 |
| 13 | 0 | 0 | 1 | 1 | $\sigma_{34}$ |
| 14 | 1 | 0 | 1 | 1 | 0 |
| 15 | 0 | 1 | 1 | 1 | 0 |
| 16 | 1 | 1 | 1 | 1 | 0 |

Table 1: Indices and values of $\boldsymbol{\sigma}^{(4)}$ when third order and higher moments are zero.

where the sum is over indices $(j_1, \ldots, j_n) \in \{0,1\}^n$. Writing out the sum over $j_n$ into two terms, we get

$$p_{k_1 \cdots k_n} = \mathbf{B}^{(n)}_{k_n 0} \sum_{j_1,\ldots,j_{n-1}} \mathbf{B}^{(n-1)}_{k_{n-1} j_{n-1}} \cdots \mathbf{B}^{(1)}_{k_1 j_1} \boldsymbol{\sigma}^{(n)}_{j_1 \cdots j_{n-1} 0} + \mathbf{B}^{(n)}_{k_n 1} \sum_{j_1,\ldots,j_{n-1}} \mathbf{B}^{(n-1)}_{k_{n-1} j_{n-1}} \cdots \mathbf{B}^{(1)}_{k_1 j_1} \boldsymbol{\sigma}^{(n)}_{j_1 \cdots j_{n-1} 1}. \quad (6)$$

Using (2) and (1), we have $\boldsymbol{\sigma}^{(n)}_{j_1 \cdots j_{n-1} 0} = \boldsymbol{\sigma}^{(n-1)}_{j_1 \cdots j_{n-1}}$ and $\boldsymbol{\sigma}^{(n)}_{j_1 \cdots j_{n-1} 1} = \boldsymbol{\gamma}^{(n-1)}_{j_1 \cdots j_{n-1}}$. It follows that the first term in (6) is $\mathbf{B}^{(n)}_{k_n 0} p_{k_1 \cdots k_{n-1}}$. We therefore have

$$p_{k_1 \cdots k_n} = \mathbf{B}^{(n)}_{k_n 0} p_{k_1 \cdots k_{n-1}} + \mathbf{B}^{(n)}_{k_n 1} \sum_{j_1,\ldots,j_{n-1}} \mathbf{B}^{(n-1)}_{k_{n-1} j_{n-1}} \cdots \mathbf{B}^{(1)}_{k_1 j_1} \boldsymbol{\gamma}^{(n-1)}_{j_1 \cdots j_{n-1}}.$$

Dividing by $p_{k_1 \cdots k_{n-1}}$ yields (5). □

For $n \in [m]$, assume $p_{k_1 \cdots k_n} > 0$ and let $\mathbf{v}^{(n)} \in \mathbb{R}^n$ be a vector defined elementwise via

$$\mathbf{v}^{(n)}_i \stackrel{\text{def}}{=} \frac{\mathbf{B}^{(i)}_{k_i 1}}{p_{k_1 \cdots k_n}} \prod_{j \in [n] \setminus \{i\}} \mathbf{B}^{(j)}_{k_j 0}. \quad (7)$$

**Proposition S1.3.** *Assume $p_{k_1 \cdots k_n} > 0$. The conditional probability $p_{k_n | k_1 \cdots k_{n-1}}$ satisfies*

$$p_{k_n | k_1 \cdots k_{n-1}} = \mathbf{B}^{(n)}_{k_n 0} + \mathbf{B}^{(n)}_{k_n 1} \mathbf{v}^{(n-1)\top} \tilde{\boldsymbol{\gamma}}^{(n-1)}. \quad (8)$$

*Proof.* Consider the sum in (5) in Lemma S1.2. An element $\boldsymbol{\gamma}^{(n-1)}_{j_1 \cdots j_{n-1}}$ in that sum can be nonzero only if $j_1 + \cdots + j_{n-1} = 1$, in which case it is a second order moment $\sigma_{in}$ with $i \in [n-1]$. Using that fact together with the definition of $\mathbf{v}^{(n-1)}$ and $\tilde{\boldsymbol{\gamma}}^{(n-1)}$ in (7) and (3), respectively, we arrive at (8). □

Our goal is to sample $X_1, \ldots, X_m$ iteratively: We first sample $X_1$, then $X_2$ given the outcome of $X_1$, then $X_3$ given the outcome of $X_1$ and $X_2$, and so forth. By leveraging Proposition S1.3, we can construct an algorithm which efficiently samples $m$ random variables from the MVB. Notice that computing an element $\mathbf{v}_i^{(n)}$ via (7) costs $O(n)$, which would bring the total cost of computing all elements of $\mathbf{v}^{(n)}$ to $O(n^2)$. Since a different $\mathbf{v}^{(n)}$ is needed for each conditional probability in (8), such an approach would bring the total cost of the sampling algorithm to $O(m^3)$. A key insight to making the sampling efficient is that $\mathbf{v}^{(n)}$ can be computed from $\mathbf{v}^{(n-1)}$ via an update which only costs $O(n)$. For $n \in [m]$, assume that $p_{k_1 \cdots k_n} > 0$ and define the constants

$$c^{(n)} \stackrel{\text{def}}{=} \frac{\mathbf{B}_{k_1 0}^{(1)} \mathbf{B}_{k_2 0}^{(2)} \cdots \mathbf{B}_{k_n 0}^{(n)}}{p_{k_1 \cdots k_n}}. \tag{9}$$

**Proposition S1.4.** *For $n \in \{2, 3, \ldots, m\}$, assuming $p_{k_1 \cdots k_n} > 0$, $\mathbf{v}^{(n)}$ and $c^{(n)}$ satisfy the recursions*

$$\mathbf{v}^{(n)} = \begin{bmatrix} \frac{\mathbf{B}_{k_n 0}^{(n)}}{p_{k_n | k_1 \cdots k_{n-1}}} \mathbf{v}^{(n-1)} \\ \frac{\mathbf{B}_{k_n 1}^{(n)}}{p_{k_n | k_1 \cdots k_{n-1}}} c^{(n-1)} \end{bmatrix} \tag{10}$$

*and*

$$c^{(n)} = \frac{\mathbf{B}_{k_n 0}^{(n)}}{p_{k_n | k_1 \cdots k_{n-1}}} c^{(n-1)}. \tag{11}$$

*Proof.* Since $p_{k_1 \cdots k_n} > 0$, this implies that $p_{k_n | k_1 \cdots k_{n-1}} > 0$, which means the divisions in (10) and (11) are well-defined. Equation (10) follows from the definition of $\mathbf{v}^{(n)}$ in (7) and the fact that $p_{k_n | k_1 \cdots k_{n-1}} = p_{k_1 \cdots k_n} / p_{k_1 \cdots k_{n-1}}$. Equation (11) follows immediately from the definition of $c^{(n)}$ in (9). □

Suppose the second order moments are stored in a matrix $\mathbf{C} \in \mathbb{R}^{m \times m}$:

$$\mathbf{C} = \begin{bmatrix} \sigma_{11} & \sigma_{12} & \sigma_{13} & \cdots & \sigma_{1m} \\ \sigma_{21} & \sigma_{22} & \sigma_{23} & \cdots & \sigma_{2m} \\ \vdots & \vdots & \vdots & & \vdots \\ \sigma_{m1} & \sigma_{m2} & \sigma_{m3} & \cdots & \sigma_{mm} \end{bmatrix}.$$

Notice that the vectors $\tilde{\boldsymbol{\gamma}}^{(n)} = \mathbf{C}_{1:n,n+1}$ can be read directly from $\mathbf{C}$.

In Algorithm S1, we give a high-level algorithm for our sampling approach. Both of our proposed methods `rb_unstr` and `rb_dplr` follow this high-level algorithm, but differ on how they perform the substeps. The steps that differ between the algorithms are the initialization on line 4, the computation of the conditional probability on line 6, and the update on line 10. In Algorithm S2 we specify how these steps are done in `rb_unstr` for a generic second moment matrix with the third and higher moments equal to zero. Notice that the division on line 4a in Algorithm S2 is well-defined, since $p_{k_1}$ must be positive. Similarly, the computations on lines 10a and 10b in Algorithm S2 involve divisions by $p_{k_n | k_1 \cdots k_{n-1}}$ which is also guaranteed to be positive. The correctness of line 6a in Algorithm S2 follows from Proposition S1.3.

**Remark S1.5** (Complexity of Algorithm S2)**.** Lines 4a and 4b both cost $O(1)$ each. Line 6a has cost $O(n)$. Lines 10a and 10b have cost $O(n)$ and $O(1)$, respectively. Since lines 6 and 10 are repeated $O(m)$ times, it follows that the total cost of the algorithm is $O(m^2)$. For a general $\mathbf{C}$ this is also optimal asymptotically, since there are up to $m(m-1)/2 \sim m^2$ nonzero elements above (or below) the diagonal in $\mathbf{C}$ that we need to consider.

**Algorithm S1:** High level algorithm for MVB sampling

**Data:** Means $\mu_1, \dots, \mu_m$ and second order moments in $\tilde{\boldsymbol{\gamma}}^{(1)}, \dots, \tilde{\boldsymbol{\gamma}}^{(m-1)}$
**Result:** One realization $(k_1, \dots, k_m)$ from MVB

```
/* Initialize: Draw X_1 */
1  Set p_1 = μ_1
2  Draw k_1 = Bernoulli(p_1)
3  if k_1 = 0 then set p_{k_1} = 1 − p_1
4  Initialize parameters
/* Recurse: Draw X_2, ..., X_m */
5  for n = 2, 3, ..., m do
6      Compute p_{1|k_1···k_{n−1}} ≝ ℙ(X_n = 1 | X_1 = k_1, ..., X_{n−1} = k_{n−1})
7      Draw k_n = Bernoulli(p_{1|k_1···k_{n−1}})
8      if k_n = 0 then set p_{k_n|k_1···k_{n−1}} = 1 − p_{1|k_1···k_{n−1}}
9      if n < m then
10         Update parameters
11     end
12 end
13 return realization (k_1, ..., k_m)
```

#### S1.1 Efficient algorithm for diagonal plus low-rank second moments

We now consider the case when $\mathbf{C} = \mathbf{D} + \mathbf{U}\mathbf{U}^\top$, where $\mathbf{D} \in \mathbb{R}^{m\times m}$ is diagonal and $\mathbf{U} \in \mathbb{R}^{m\times r}$ for $r < m$. As discussed in Remark S1.5, the main cost in Algorithm S2 are on lines 6a and 10a. When $\mathbf{C}$ is diagonal-plus-low-rank, there is a clear relationship between subsequent vectors $\tilde{\boldsymbol{\gamma}}^{(n)}$. This can be leveraged to avoid computing the inner product on line 6a. Moreover, the step on line 10a can also be avoided: Instead of computing $\mathbf{v}^{(n)}$, we can instead keep track of $r$ inner products that are fast to update from one step of the for loop to another.

Let $\mathbf{u}^{(\ell)}$ be the $\ell$th column of $\mathbf{U}$. Then

$$\mathbf{C} = \mathbf{D} + \sum_{\ell=1}^{r} \mathbf{u}^{(\ell)}\mathbf{u}^{(\ell)\top} = \mathbf{D} + \begin{bmatrix} \sum_{\ell=1}^r \mathbf{u}^{(\ell)}\mathbf{u}_1^{(\ell)} & \sum_{\ell=1}^r \mathbf{u}^{(\ell)}\mathbf{u}_2^{(\ell)} & \cdots & \sum_{\ell=1}^r \mathbf{u}^{(\ell)}\mathbf{u}_m^{(\ell)} \end{bmatrix}.$$

**Algorithm S2:** Generate Bahadur order-2 MVB variates with arbitrary correlation structures (`rb_unstr`)

```
/* Same as Algorithm S1 with lines 4, 6 and 10 defined as below */
4   Initialize parameters:
4a      v^(1) = B^(1)_{k_1 1} / p_{k_1}
4b      c^(1) = 1
6   Compute p_{1|k_1···k_{n−1}}:
6a      p_{1|k_1···k_{n−1}} = μ_n + v^(n−1)⊤ γ̃^(n−1)
10  Update parameters:
10a     Compute v^(n) according to (10)
10b     Compute c^(n) according to (11)
```

It follows that

$$\tilde{\boldsymbol{\gamma}}^{(n)} = \sum_{\ell=1}^{r} \mathbf{u}_{1:n}^{(\ell)} \mathbf{u}_{n+1}^{(\ell)}, \qquad n \in [m-1].$$

The inner product $\mathbf{v}^{(n)\top}\tilde{\boldsymbol{\gamma}}^{(n)}$ can be written

$$\mathbf{v}^{(n)\top}\tilde{\boldsymbol{\gamma}}^{(n)} = \sum_{\ell=1}^{r} \mathbf{v}^{(n)\top}\mathbf{u}_{1:n}^{(\ell)} \mathbf{u}_{n+1}^{(\ell)} = \sum_{\ell=1}^{r} x_\ell^{(n)} \mathbf{u}_{n+1}^{(\ell)},$$

where each $x_\ell^{(n)} \overset{\text{def}}{=} \mathbf{v}^{(n)\top}\mathbf{u}_{1:n}^{(\ell)}$ is an inner product. As Proposition S1.6 shows, these $r$ numbers can be updated efficiently.

**Proposition S1.6.** *Assume $p_{k_n|k_1\cdots k_{n-1}} > 0$. For each $\ell \in [r]$ and $n \in \{2, 3, \ldots, m-1\}$, the numbers $x_\ell^{(n)}$ satisfy the recursion*

$$x_\ell^{(n)} = x_\ell^{(n-1)} \frac{\mathbf{B}_{k_n 0}^{(n)}}{p_{k_n|k_1\cdots k_{n-1}}} + c^{(n-1)} \frac{\mathbf{B}_{k_n 1}^{(n)} \mathbf{u}_n^{(\ell)}}{p_{k_n|k_1\cdots k_{n-1}}}, \tag{12}$$

*where $c^{(n-1)}$ is defined as in (9).*

*Proof.* Using (10),

$$\begin{aligned} x_\ell^{(n)} &= \mathbf{v}^{(n)\top}\mathbf{u}_{1:n}^{(\ell)} \\ &= \begin{bmatrix} \mathbf{B}_{k_n 0}^{(n)} (p_{k_n|k_1\cdots k_{n-1}})^{-1} \mathbf{v}^{(n-1)\top} & \mathbf{B}_{k_n 1}^{(n)} (p_{k_n|k_1\cdots k_{n-1}})^{-1} c^{(n-1)} \end{bmatrix} \begin{bmatrix} \mathbf{u}_{1:n-1}^{(\ell)} \\ \mathbf{u}_n^{(\ell)} \end{bmatrix} \\ &= \mathbf{B}_{k_n 0}^{(n)} (p_{k_n|k_1\cdots k_{n-1}})^{-1} \mathbf{v}^{(n-1)\top}\mathbf{u}_{1:n-1}^{(\ell)} + \mathbf{B}_{k_n 1}^{(n)} (p_{k_n|k_1\cdots k_{n-1}})^{-1} c^{(n-1)} \mathbf{u}_n^{(\ell)}. \end{aligned}$$

Recognizing that $\mathbf{v}^{(n-1)\top}\mathbf{u}_{1:n-1}^{(\ell)} = x_\ell^{(n-1)}$ and rearranging, we arrive at (12). □

Algorithm S3 shows how `rb_dplr` is implemented. It uses Proposition S1.6 to reduce the complexity of Algorithm S2 in the case when $\mathbf{C}$ is diagonal-plus-low-rank. Notice that the lines 4a and 10a of Algorithm S3 involve divisions by $p_{k_1}$ and $p_{k_n|k_1\cdots k_{n-1}}$, respectively, which are both guaranteed to be positive, so these computations are well-defined.

---

**Algorithm S3:** Generate Bahadur order-2 MVB variates with diagonal-plus-low-rank (DPLR) correlation structures (`rb_dplr`)

---

`/* Same as Algorithm S1 with lines 4, 6 and 10 defined as below */`

**4** Initialize parameters:

**4a**    **for** $\ell \in [r]$ **do** set $x_\ell^{(1)} = \mathbf{B}_{k_1 1}^{(1)} \mathbf{u}_1^{(\ell)} / p_{k_1}$

**4b**    $c^{(1)} = 1$

**6** Compute $p_{1|k_1\cdots k_{n-1}}$:

**6a**    $p_{1|k_1\cdots k_{n-1}} = \mu_n + \sum_{\ell=1}^{r} x_\ell^{(n-1)} \mathbf{u}_n^{(\ell)}$

**10** Update parameters:

**10a**    **for** $\ell \in [r]$ **do** compute $x_\ell^{(n)}$ according to (12)

**10b**    Compute $c^{(n)}$ according to (11)

---

**Remark S1.7** (Complexity of Algorithm S3)**.** Lines 4a and 4b cost $O(r)$ and $O(1)$, respectively. Line 6a costs $O(r)$. Lines 10a and 10b cost $O(r)$ and $O(1)$, respectively. Since lines 6 and 10 are repeated $O(m)$ times, it follows that the total cost of the algorithm is $O(mr)$. For a general $\mathbf{U}$ this is optimal asymptotically, since there are up to $mr$ nonzero elements in $\mathbf{U}$ that need to be considered.

**Remark S1.8** (Exchangeable correlation)**.** With exchangeable correlation structure, all off-diagonal elements of $\mathbf{C}$ are the same, i.e., $\mathbf{C}_{ij} = \alpha$ when $i \neq j$. This is a diagonal-plus-rank-1 structure: With $\mathbf{u} \in \mathbb{R}^m$ such that all entries equal to $\sqrt{\alpha}$, $\mathbf{C} = \mathbf{D} + \mathbf{u}\mathbf{u}^\top$ for some diagonal matrix $\mathbf{D}$. Algorithm S3 therefore handles this case with a complexity of $O(m)$.

### S2 Using MVB random variates to model AM

#### S2.1 Equilibrium distribution of causal variants under AM

We consider the equilibrium distribution of haploid causal variants $X_1, \ldots, X_m$ with allele frequencies $\mu_1, \ldots, \mu_m$ under primary-phenotypic assortative mating for an additive phenotype with panmictic heritability $h_0^2$, panmictic genetic variance $\sigma_{g,0}^2 = h_0^2$, and cross-mate phenotype correlation $r$. Following [2], the equilibrium heritability is

$$h_\infty^2 = \frac{1}{2r}\left((1-h_0^2)^{-1} - \sqrt{(1-h_0^2)^{-2} - 4rh_0^2(1-h_0^2)^{-1}}\right),$$

and the equilibrium cross-mate genetic correlation and genetic variance are respectively $r_{g,\infty} = r \cdot h_\infty^2$ and $\sigma_{g,\infty}^2 = \sigma_{g,0}^2/(1-r_{g,\infty})$. Additionally, we denote the equilibrium phenotypic variance $\sigma_{y,\infty}^2$ and the standardized haploid effects $\boldsymbol{\beta}$.

Assuming casual variants are unlinked at panmixis, the correlation matrix of causal haploid variants will be of the form $\mathbf{R} = \mathbf{D} + \boldsymbol{\phi}\boldsymbol{\phi}^T$ where, following [3], $\boldsymbol{\phi} : \mathbb{R}^m \to \mathbb{R}^m$ is a function of the standardized haploid effects $\boldsymbol{\beta}$ given elementwise by

$$\phi_k = \sqrt{\sigma_{y,\infty}^2 \mu_k(1-\mu_k)/(8\beta_k^2 r)}\left(\sqrt{4\beta_k^2 r/\sigma_{y,\infty}^2 + (1-r_{g,\infty})^2} - (1-r_{g,\infty})\right), \tag{13}$$

and $\mathbf{D}$ is the diagonal matrix with entries $\mathbf{D}_{kk} = 1 - \phi_k^2$. Setting $\mathbf{U} = \boldsymbol{\phi}$ allows drawing haploid causal variants from the equilibrium distribution under AM via `rb_dplr`.

The `rBahadur` library includes several utility functions for simulating genotypes under this model. `vg_eq`, `h2_eq`, and `rg_eq` compute equilibrium parameters given initial conditions. `am_covariance_structure` computes $\boldsymbol{\phi}$ via (13). Finally, `am_simulate` automates the entire simulation procedure, taking initial parameters as inputs and returning genotypes, effects, and phenotypes.

#### S2.2 Generating genotypes under AM with complex local LD

The greatest limitation of using the proposed methods to simulate genotype data is that the desired first and second moments must result in a valid Bahadur order-2 MVB distribution. Specifically, from Proposition 3 of [4], we must have that the minimal eigenvalue of the target correlation matrix $\mathbf{R}$ satisfies

$$\lambda_{\min}(\mathbf{R}) \geq 1 - 2/\sum_{j=1}^m \max\left\{\frac{\sigma_j}{1-\sigma_j}, \frac{1-\sigma_j}{\sigma_j}\right\},$$

where $\sigma_j = \sqrt{\mu_j(1-\mu_j)}$. In practice, this makes it difficult to accommodate strong local LD among nearby loci. To address this limitation, we propose the two step procedure outlined in Algorithm S4. Specifically, we first generate causal variants using `rb_dplr` and empirical allele frequencies at $m$ of $p \gg m$ SNPs across the genome. We then determine a partition of the genome such that each slice contains one and only one causal locus, using a recombination map to select boundaries. For each resulting haplotype block, we then choose haplotypes from a real or synthetic panmictic founder population from among those which agree with the simulated variant at the casual locus. Proceeding in this manner, we are able to benefit from the computational efficiency of `rb_dplr` while generating genotype data with realistic local LD.

**Algorithm S4:** High-level algorithm for generating synthetic genotypes under AM while preserving complex LD structure

**Data:** $N$ founder's haploid variants at $p$ diploid loci $\{h_{1j}\}_{j=1}^{p}, \ldots \{h_{Nj}\}_{j=1}^{p}$, recombination probabilities $r_2, \ldots, r_p$, causal variant indices $\{i_1, \ldots, i_m\} \subset \{1, \ldots, p\}$ for $m \ll p$ with corresponding haploid allele frequencies $\mu_1, \ldots, \mu_m$ and haploid causal effects $\beta_1, \ldots, \beta_m$, and phenotypic mate correlation $r$

**Result:** One sample of haploid variants $(X_1, \ldots, X_p)$

/* Draw MVB random variates corresponding to haploid causal variants */

Construct DPLR covariance structure parameter $\boldsymbol{\phi}$ via (13)

Draw MVB haploid causal variants $(X_{i_1}, \ldots, X_{i_m})$ using `rb_dplr` with $\mathbf{U} = \boldsymbol{\phi}$

/* Set boundaries for $m$ contiguous haplotype blocks that partition $\{1, \ldots, p\}$ */

Set first and last haplotype boundaries $b_1 = 1$, $b_{m+1} = p$

**for** $k = 2, \ldots, m$ **do**

  Rescale recombination probabilities $\mathbf{r}_k = (r_{i_{k-1}+1}, \ldots, r_{i_k})/(r_{i_{k-1}+1} + \cdots + r_{i_k})$

  Randomly select boundary $b_k \in \{i_{k-1}+1, \ldots, i_k\}$ from the corresponding categorical distribution with probabilities $\mathbf{r}_k$

**end**

/* Populate loci $\{1, \ldots, p\} \setminus \{i_1, \ldots, i_m\}$ by sampling founder haplotype blocks matching causal variants */

**for** $k = 1, \ldots, m$ **do**

  Draw founder index $n_k = \text{Uniform}(\{n \in \{1, \ldots, N\} : h_{ni_k} = X_{i_k}\})$

  /* Populate loci surrounding causal variant */

  **for** $j = b_{k-1}, \ldots, b_{k+1}$ **do**

    Set $X_j = h_{n_k j}$

  **end**

**end**

**return** haplotypes $(X_1, \ldots, X_p)$
